# Finding metabolic pathways in large networks through atom-conserving substrate-product pairs

**DOI:** 10.1101/2020.11.25.398453

**Authors:** Jasmin Hafner, Vassily Hatzimanikatis

## Abstract

Finding biosynthetic pathways is essential for metabolic engineering of organisms to produce chemicals, biodegradation prediction of pollutants and drugs, and for the elucidation of bioproduction pathways of secondary metabolites. A key step in biosynthetic pathway design is the extraction of novel metabolic pathways from big networks that integrate known biological, as well as novel, predicted biotransformations. However, especially with the integration of big data, the efficient analysis and navigation of metabolic networks remains a challenge. Here, we propose the construction of searchable graph representations of metabolic networks. Éach reaction is decomposed into pairs of reactants and products, and each pair is assigned a weight, which is calculated from the number of conserved atoms between the reactant and the product molecule. We test our method on a biochemical network that spans 6,546 known enzymatic reactions to show how our approach elegantly extracts biologically relevant metabolic pathways from biochemical networks, and how the proposed network structure enables the application of efficient graph search algorithms that improve navigation and pathway identification in big metabolic networks. The weighted reactant-product pairs of an example network and the corresponding graph search algorithm are available online. The proposed method extracts metabolic pathways fast and reliably from big biochemical networks, which is inherently important for all applications involving the engineering of metabolic networks.

## Introduction

Extracting meaningful metabolic pathways from large metabolic networks is essential for the computational design of bioproduction pathways, for the elucidation of biosynthesis of natural products, and for the fundamental understanding of metabolism. Metabolic pathways, which describe the transformation from one or several source molecules over consecutive reaction steps into one or several target molecules, provide the roadmap guiding these various applications^1–3^. Traditionally, metabolic pathways were drawn by hand after directly inferring biochemical transformations from experimental evidence. However, the advent of the omics era and the dramatic increase of computational data resources has drastically changed the way we study biochemistry. Our knowledge is now collected in continuously growing databases, providing new opportunities for fundamental research and metabolic engineering, making the by-hand design of pathways nearly impossible and obsolete. Additionally, these new resources can be used to design non-canonical pathways that do not exist in nature. While many non-natural pathways have been historically designed by intuition using paper and pencil, it is likely that more efficient solutions will be missed. To address this challenge, computational pathway search tools have been developed to extract metabolic pathways from biochemical databases^4,5^.

Overall, a biosynthesis pathway converts one or several metabolites into a final target metabolite containing all or most of the atoms found in the precursor compound or converts a more complex metabolite back into simpler building blocks that conserve the atoms of the original metabolite. Hence, unless stated otherwise, atom conservation between start and end point defines a metabolic pathway. If we want to find biosynthetic or biodegradation pathways, we therefore have to look for atom-conserving reaction paths connecting start and end metabolite(s). The objective of pathway search methods is to find “biologically meaningful” pathways, which are here defined as biochemical routes that fulfill the following criteria: (i) Core atoms are conserved throughout the pathway, (ii) loops are not allowed, meaning that no metabolite appears twice, and (iii) other metabolites that contribute to the main biotransformation route in a lesser degree are considered as cofactors or co-substrates. Expressed in a more general way, we aim to recover linear metabolic pathways as they are shown in textbooks, but through an automated approach.

There are different ways to mathematically describe a metabolic network (i.e., stoichiometric matrix or mathematical graph), and hence different approaches to analyze these networks and find pathways^4^. However, optimization-based methods to retrieve pathways from stoichiometric matrices are not applicable to big biochemical networks, and therefore not further discussed here. Graph-based methods on the other hand are suitable for large-scale applications due to their computational efficiency. Different methodologies have been developed in the past to (i) represent metabolic networks as mathematical graph structures, and (ii) to find pathways within the graph from a given source to a target metabolite. Different approaches have been explored to bias a biochemically blind graph search algorithm towards biologically meaningful pathways, such as the exclusion of cofactors from the network, defining reactant pairs through the chemical similarity of compounds^6^, precomputing recurring subpaths^7^, and atom or substructure conservation throughout the pathway^6,8^. Atom conservation in general, and carbon conservation specifically, have been shown to be a valuable criterion for finding biologically meaningful pathways^9–14^.

There are several solutions to pathway discovery that employ the concept of atom conservation. Initially, the tracking of single atoms was used by Arita *et al*. to calculate network properties of the metabolism of *Escherichia coli*^9^. Later, atom tracking was used to improve the quality of pathway search tools by ensuring that one or several atoms were conserved throughout the pathway^10,11,15–18^. Atom-tracking methods have been shown to find biologically relevant pathways, although the high quality came with an increased computational cost. An alternative strategy has been pursued by the Kyoto Encyclopedia of Genes and Genomes (KEGG). Their reactions are annotated with chemical structure alignments, also called substrate-product pairs or reactant pairs (short RPAIRS). KEGG’s pathway prediction server, named PathPred, uses the reactant pairs to create a searchable graph of biologically meaningful biotransformations^19^. Instead of tracking atoms individually, PathPred approximates the atom conservation by defining atom-conserving reactant pairs, which decreases the complexity of the path search problem. However, their classification system is based on a combination of manual curation and automatic annotation, a strategy that is not easily applicable to large biochemical networks, such as the ATLAS of Biochemistry with its more than 140,000 predicted reactions^20,21^. Large biochemical databases, especially those including hypothetical reactions, require reliable and computationally efficient algorithms to extract biologically relevant biochemical pathways.

Here, we address the challenge of efficiently searching and analyzing big biochemical networks. We propose a new method, named NICEpath, that biases the graph search towards atom-conserving pathways. To achieve this, we calculate weighted reactant-product pairs that reflect the atom conservation in each reaction, and we use the atom-conserving pairs to represent biochemical reaction networks as weighted graphs that are compatible with efficient search algorithms. The pathways found by NICEpath therefore fit the definition of “biologically meaningful” in the sense that they fulfill the three criteria mentioned earlier. The algorithm finds atom-conserving pathways first and returns a pathway list ranked according to atom conservation. NICEpath can be readily employed to extract and compare metabolic pathways from overall biochemical database (e.g., KEGG) or from metabolic networks specific to an organism (e.g., genome-scale models). The method can be further applied to efficiently search large biochemical networks, as they are generated by reaction prediction tool such as BNICE.ch^22^.

## Materials and Methods

Our approach can be divided into four steps (Figure 1): (i) The first step consists of acquiring an atom-level representation of each reaction. The atom maps can come from databases, atom-mapping algorithms, or, in our case, enzymatic reaction rules as implemented in BNICE.ch^22^. (ii) In a second step, each atom-mapped reaction is decomposed into all the possible reactant-product pairs. For each pair, we calculate the Conserved Atom Ratio (CAR) from the number of conserved atoms between reactant and product and the size of the molecules in terms of number of atoms. (iii) The atom-weighted substrate-product pairs are used to construct a weighted undirected graph, where the distance between reactants and products are inversely proportional to the CAR. (iv) Once the graph of weighted substrate-product pairs is constructed, we can apply well-established graph search methods to find the shortest paths, which will inherently find the pathways that conserve the highest number of atoms. NICEpath uses the Yen’s k-shortest loop-less path^23^ algorithm, a standard method to find a predefined number (k) of shortest paths in weighted graph, avoiding the repetition of nodes.

**Figure 1:**
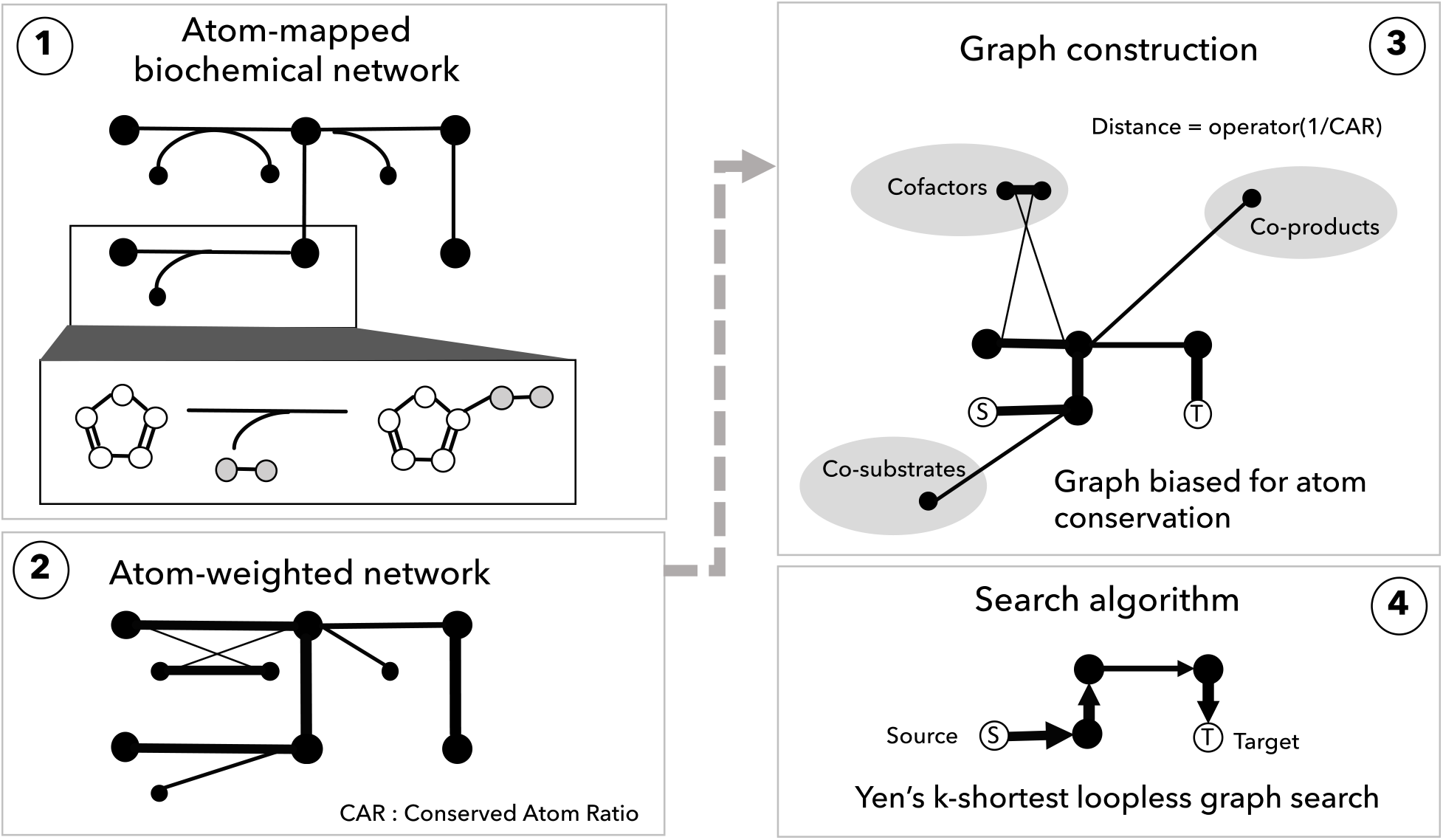
The workflow of the pathway search is divided into two parts. The first two steps (left) describe the atom weighting of the network from atom-mapped reactions. In this study, steps 1 and 2 are performed by BNICE.ch. Steps 3 and 4 (right), implemented in NICEpath, take the atom-weighted network as an input to create a searchable graph structure and finally apply a Yen’s k-shortest pathway search.

### Biochemically correct atom mapping with BNICE.ch

Atom-mapped reactions are the prerequisite for calculating weighted reactant-product pairs. Here, we use the computational tool BNICE.ch, developed to predict hypothetical biochemical networks, to calculate biochemically correct atom mappings of enzymatic reactions. The core of BNICE.ch consists of 442 bidirectional, generalized biochemical reaction rules that describe the biochemical reaction mechanisms of enzymatic reactions. The reaction rules are applied to a molecular structure to (i) reconstruct atom-mapped, known biochemical reactions; and (ii) to predict all possible biochemical transformations that a given compound can undergo along with the product compounds generated in the process. Here, BNICE.ch calculates atom maps for metabolic reactions using the mechanistic knowledge stored in the reaction rules, as described by Hadadi *et al*.^24^ In this step, other tools for the automatic atom mapping of reactions may also be applied to generate atom maps^17,18,25^.

### Calculation of weighted reactant-product pairs

The following steps are applied to each reaction in the network to generate atom-weighted reactant-product pairs: (i) Each reaction is split into all possible reactant-product pairs. (ii) For each pair of reactant and product, the number of common atoms (*n_c_*) between reactant and product is calculated along with the total number of atoms in the reactant (*n_r_*) and the total number of reactants in the product (*n_p_*). Hydrogen atoms are omitted from the calculation. (iii) For each pair, the ratio of conserved atoms (in the following, called Conserved Atom Ratio, or CAR) is calculated with respect to the reactant (*CAR_r_*) and with respect to the product (*CAR_p_*).

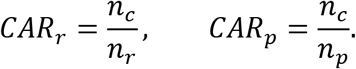

(iv) To calculate a bidirectional CAR, the mean CAR is multiplied with a correction factor that increases with the difference between the number of common atoms and the total number of atoms in the molecule.

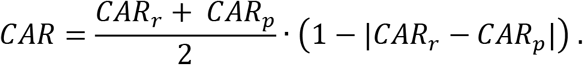

The only exception to this approach is made for reactions involving the cofactor Coenzyme A (CoA). In a molecule, CoA is treated as a single atom when it occurs in both the reactant and in the product, mainly because the high number of conserved atoms between the comparably big CoA leads to high CARs, thus masking the biochemically more interesting connections between the smaller metabolites that are attached to and detached from CoA during metabolic transformations. The final CAR value is used to weight reactant-product pairs in the network.

### Assigning mechanisms to biochemical reactions from the KEGG reference network

We used KEGG as a reference database for enzymatic reactions, from which we extracted all reactions that have an associated mechanism in BNICE.ch. If a given reaction from KEGG could be reconstructed with BNICE.ch, it was assigned a reaction mechanism that allowed us to retrieve the number of conserved atoms between each reactant-product pair. The set of KEGG reactions with assigned reaction mechanisms and pre-calculated CAR values was used for further validation and as an example network for network analysis and pathway search. The set of BNICE.ch curated KEGG reactions is available from the GitHub repository at https://github.com/EPFL-LCSB/nicepath.

### Graph representation of biochemical networks

For a given reaction network, NICEpath loads all the reactant-product pairs to generate a weighted, undirected graph, where metabolites are nodes connected by edges, representing the reactant-product relationship. Edges are assigned a weight that defines the relation between two connected nodes. To use state-of-the-art shortest-path graph search algorithms, highly atom-conserving reactants should be close to each other, and pairs that only share a few atoms should be further away. Hence, we convert the CAR into a distance:

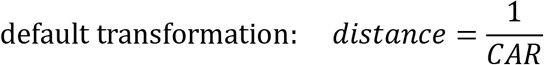

NICEpath accepts two alternative ways to calculate the distance, which can be used to modulate the influence of the atom conservation on the weight of the reactant-product pair.

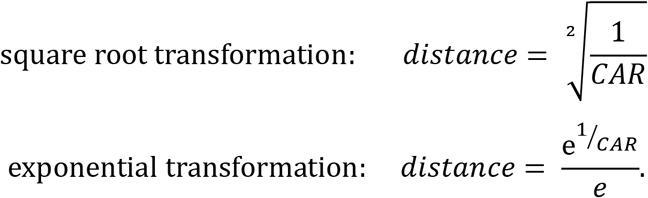

The type of transformation can be changed to square root or exponential depending on the nature of the pathway search problem, i.e., the structures of source and target molecules as well as the estimated number of biotransformation used to convert one into the other. The distance measure is used to reconstruct a directed graph whose edge weights represent the atomic distance between reactants and products. For longer pathways, we recommend using the exponential transformation because it increases the penalty for pairs with low CARs, which makes the search more conservative in terms of atoms.

For this study, we grouped duplicate KEGG compounds into one node. Duplicates were identified based on the first fourteen letters of the InChIKey, which means that different stereoisomers of the same molecular structure were merged into one node.

### Finding metabolic pathways with graph search

NICEpath applies a Yen’s k-shortest loop-less path search^23^ to extract the shortest pathways from the weighted network of reactant-product pairs using the python package NetworkX. As inputs, the pathway search algorithm takes a weighted graph, a source compound, a target compound, and the maximum number of shortest paths (k) to be found. As soon as this number k is reached, the algorithm stops and returns all the k-shortest paths in terms of summed edge weights.

The run time of NICEpath depends on the structure of the network, the distance between the source and target compound in the graph, the number of pathways to be found, and the maximum pathway length allowed. As an example, to find 10,000 pathways of maximum length 100, the algorithm runs for about 15 minutes on a standard desktop computer using a single core. If there are several source compounds given as input, NICEpath can run path searches in parallel for different source compounds using all available cores.

### Network analysis in NICEpath

NICEpath first calculates standard network statistics, such as the number of nodes and edges, and then extracts an undirected, unweighted network from the original network by only considering edges with a CAR higher than a given threshold. For this new network, the number of components, or disjoint graphs, is extracted, and the biggest component is further analyzed regarding its size relative to the previous network as well as its diameter. Since searching for pathways between two compounds belonging to different disconnected graphs will not yield any good pathways, NICEpath will warn the user in this case.

### Software

The NICEpath code can be executed with any python version up to 3.7. The NetworkX python library (https://networkx.github.io/) was used to implement and search the reaction graph. An extensive list of libraries used can be found in the specification file on GitHub.

## Results and Discussion

### Weighted substrate-product pairs capture the main biotransformations

To validate the biochemical relevance of weighted substrate-product pairs, we compared them to the KEGG RPAIR database consisting of manually curated, atom-mapped, substructure-conserving reactant-product pairs, called RPAIRs^26^. KEGG differentiates between five types of RPAIRS: “main”, “cofac”, “trans”, “ligase”, and “leave”. The four latter ones describe cofactor pairs, small groups transferred by transferases, nucleotide triphosphate consumption by ligases, and the addition or removal of small inorganic compounds by lyases and hydrolases, respectively. The first type, “main”, describes the main biotransformation in a given reactions. To take the alcohol dehydrogenase reaction as an example, the main pair would be the transformation of the primary alcohol to the aldehyde, and the conversion of the cofactor NAD+ to NADH would be of type “cofac” (Figure 2). “Main” pairs that are used to draw the KEGG metabolic pathway maps. Therefore, a method that accurately predicts KEGG RPAIRS of type “main” can be used to reconstruct biologically relevant metabolic pathways. It should be noted that KEGG discontinued the manual definition and curation of RPAIRS in 2016, and replaced the concept of RPAIRS with an automatically calculated alternative, RCLASS.

**Figure 2:**
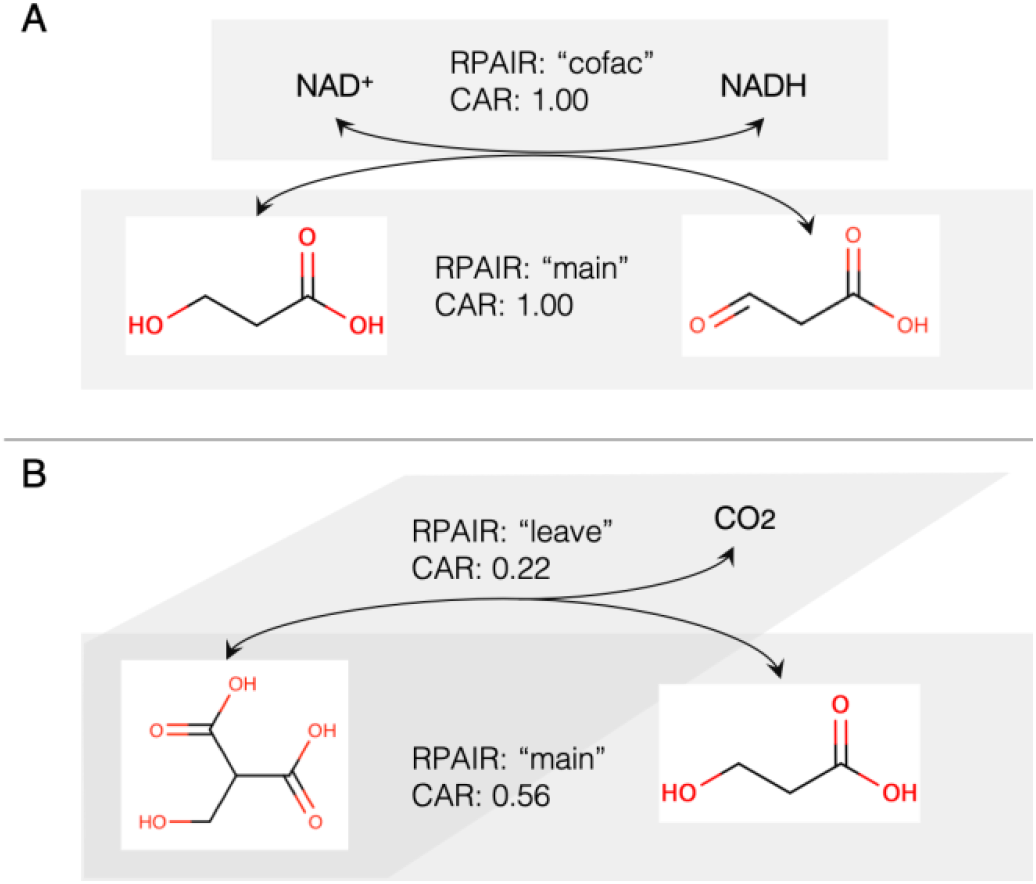
Example of relation between KEGG RPAIRs and the CAR value in a biochemical reaction. (A) Alcohol dehydrogenase, (B) decarboxylation reaction.

We validated the NICEpath method by predicting KEGG RPAIRS of type “main” using the concept of the Conserved Atom Ratio. We used BNICE.ch to calculate CAR values for a test set of 6,546 KEGG reactions for which the exact reaction mechanism is known, and which are, therefore, reconstructed by BNICE.ch (Supplementary Table S1). From these 6,546 reactions, we determined 10,747 substrate-product pairs with a non-zero CAR, meaning that at least one nonhydrogen atom is conserved between the substrate and the product (Supplementary Table S2). Out of these 10,747 pairs, 5,148 were found to be KEGG RPAIRS of type “main”. Since RPAIRs are defined based on the conservation of structural moieties within a reaction, we hypothesized that the higher the CAR value, the more atoms conserved between a substrate and a product, and hence the higher the probability that the pair would be a KEGG RPAIR of type “main”. We should therefore be able to predict the membership of a pair to the set of “main” KEGG RPAIRS by using a given CAR threshold as a classifier.

To test our hypothesis that the CAR is a good predictor for a reactant-product pair to be of KEGG RPAIR type “main”, we performed a Receiver-Operator Characteristic (ROC) analysis (Figure 3). The reference for true pairs were the 5,148 “main” RPAIRs (true positives), and the remaining 5,599 pairs were true negatives. For 100 CAR cutoff values between zero and one we calculated the number of good predictions (i.e., number of pairs with a CAR above the cutoff and of type “main”, or true positives) and bad predictions (i.e., number of pairs with a CAR above the cutoff and not of type “main”, or false positives). By drawing true positives versus false positives, we found an Area Under Curve (AUC) of 0.88. An AUC above 0.8 is considered an “excellent discrimination”^27^. We further show the tradeoff between sensitivity and specificity, as well as the Youden’s index (i.e., sensitivity + specificity - 1) to characterize this tradeoff^28^ and to determine an optimal CAR cutoff. We found that the Youden’s index is maximal at a CAR equal to 0.34, which suggests that this is the optimal CAR cutoff to tell whether a given substrate-product pair conserves enough atoms to be considered a “main” pair. This analysis shows that we can reliably use the CAR to predict KEGG RPAIRS of type “main”. The network of weighted KEGG reactant pairs for 6,546 KEGG reactions is included in the NICEpath software and used as a reaction database in the default search.

**Figure 3:**
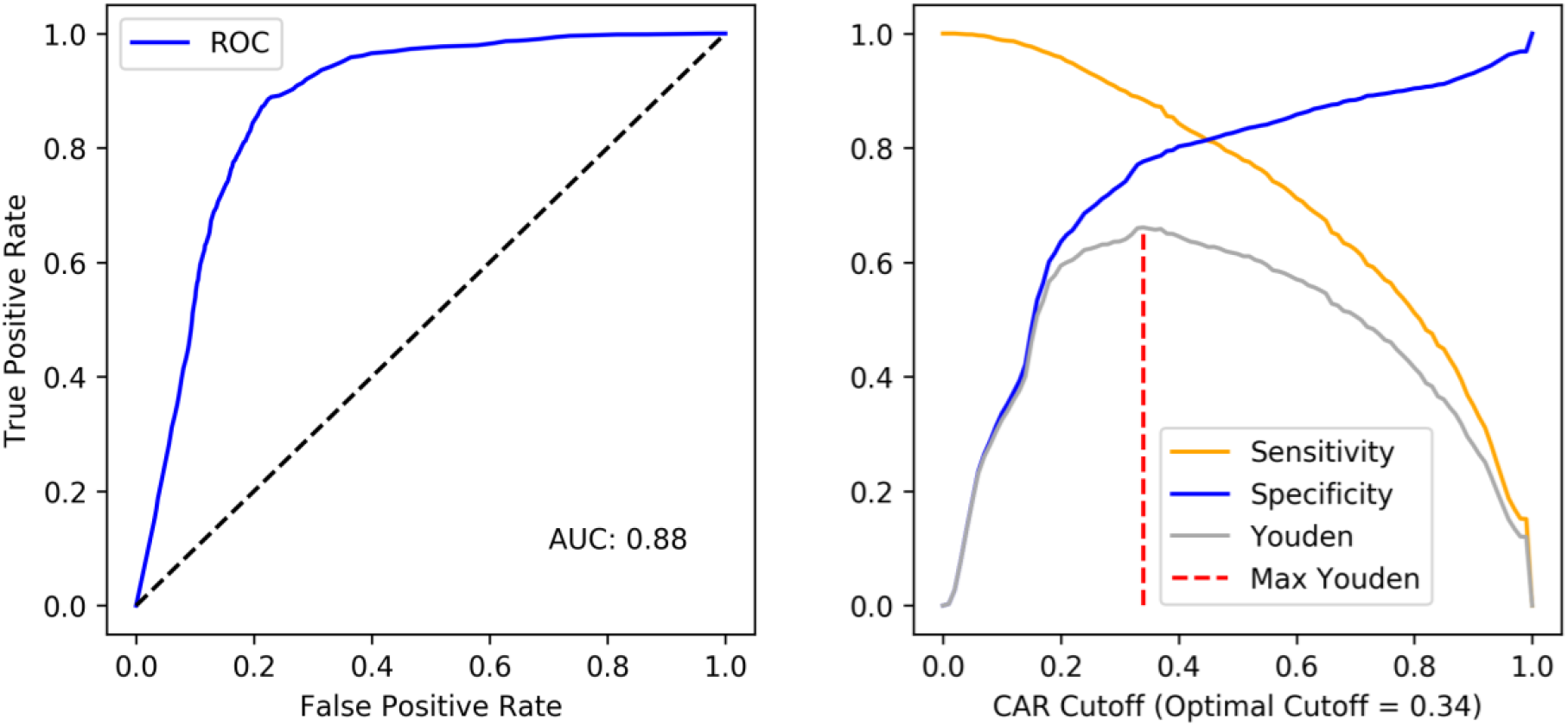
The ROC **(left panel)** curve shows the prediction of KEGG RPAIRS of type “main” by CAR score from BNICE.ch. The **right panel** shows the tradeoff between specificity (blue line) and sensitivity (yellow line). The Youden’s index (grey line) reaches its maximum (0.66) at a CAR value of 0.34 (red dashed line).

### Graph-theoretical analysis of metabolic networks

Characterizing biochemical networks from a graph-theoretical point of view can be used to evaluate the quality and connectivity of the represented network, and also bring new insights into the overall organization of metabolism. Furthermore, knowing the graph-theoretical properties of a biochemical network can be crucial for anticipating potential problems in the pathway search. NICEpath provides basic network statistics that allow us to assess the quality of the data. Here, the weighted graph of the KEGG network used for validation initially contained 5,578 compounds, or nodes, and 20,911 directed edges representing reactant-product pairs.

Certain graph properties are not defined for weighted directed graphs, such as the number of components or the network diameter. For calculating these properties, a simple, non-directed graph was generated by removing reactant-product pairs with a CAR lower than the previously calculated optimal threshold of 0.34 and by removing the weights on the remaining reactant-product pairs. The unweighted graph contains 5,518 nodes and 5,541 edges, which are distributed over 813 smaller disjoint graphs, or so-called components. The biggest component contains 2,663 nodes (48%) and 3,422 edges (62%), and it has a network diameter of 40. In other words, the longest shortest pathway connecting two compounds counts 40 biotransformation steps in the main component of the KEGG network. This means that our KEGG network is dominated by one big component, or “island”, that includes half of the metabolites in KEGG and represents the core metabolism plus connected secondary metabolism. The remaining metabolites are organized in small, disconnected subnetworks, which we hypothesize to be mostly secondary metabolites without defined biosynthesis pathways.

### Finding biologically relevant pathways with NICEpath

To illustrate the output of NICEpath, we discuss two example pathway searches. In the first example, we tried to biochemically connect tyrosine to caffeate, and we allowed a maximum number of ten pathways to be found. The pathway search resulted in ten pathways with lengths ranging from two to six consecutive reaction steps (Table 1). The quality of the pathway can be estimated from the pathway score and the average CAR. The pathway score sums the distances for each reactant-product pair in the pathway. The score reflects both the length of the pathway as well as the quality of atom conservation within the pathway, and it is eventually used by NICEpath to rank the paths. The average CAR estimates the quality of the pathway by averaging the atom conservation over each reaction step.

**Table 1:** Output of example pathway search from tyrosine (C00082) to caffeate (C01197). KEGG identifiers are used to specify compounds and reactions. The maximum number of pathways (k) was set to 10, and only one reaction alternative was printed when several reactions could do the same biotransformation.

| Index | Pathway length | Intermediates | Reaction IDs | Pathway score | Average CAR |
| --- | --- | --- | --- | --- | --- |
| 1 | 2 | C00082->C00811->C01197 | R00737->R07826 | 2.24 | 0.89 |
| 2 | 4 | C00082->C00811->C00223->C00323->C01197 | R00737->R01616->R07436->R01943 | 4.32 | 0.93 |
| 3 | 4 | C00082->C01179->C03672->C00811->C01197 | R00729->R03336->R08766->R07826 | 4.33 | 0.93 |
| 4 | 4 | C00082->C00079->C00423->C00811->C01197 | R07211->R00697->R02253->R07826 | 4.52 | 0.89 |
| 5 | 5 | C00082->C00826->C00079->C00423->C00811->C01197 | R00732->R00691->R00697->R02253->R07826 | 6.27 | 0.81 |
| 6 | 6 | C00082->C01179->C03672->C00811->C00223->C00323->C01197 | R00729->R03336->R08766->R01616->R07436->R01943 | 6.41 | 0.94 |
| 7 | 6 | C00082->C00811->C00223->C00323->C00406->C01494->C01197 | R00737->R01616->R07436->R01942->R02194->R03366 | 6.44 | 0.93 |
| 8 | 6 | C00082->C00079->C00423->C00540->C00223->C00323->C01197 | R07211->R00697->R02255->R08815->R07436->R01943 | 6.50 | 0.93 |
| 9 | 6 | C00082->C00811->C00423->C00540->C00223->C00323->C01197 | R00737->R02253->R02255->R08815->R07436->R01943 | 6.50 | 0.93 |
| 10 | 6 | C00082->C00079->C00423->C00811->C00223->C00323->C01197 | R07211->R00697->R02253->R01616->R07436->R01943 | 6.60 | 0.91 |

Out of these ten best pathways, the pathways ranked first, second, and fifth were chosen for visual inspection (Figure 4). The first pathway had a very low score of 2.24 combined with a high average CAR (0.89) and a length of two, which indicates that the pathway is of good quality because it only requires a small number of steps, all of them showing high atom conservation. Indeed, KEGG proposes this pathway in its phenylpropanoid biosynthesis map, meaning that it is biologically relevant. The second pathway, although longer, has a similarly high average CAR of 0.93, a length of four steps, and it can also be found in KEGG. To contrast these two good pathway examples with a poor example, the pathway ranked fifth shows a slightly lower average CAR of 0.81, which is due to the attachment and subsequent detachment of a one-carbon unit. In this pathway, out of five reaction steps, the last step is redundant with the first pathway, while the first four steps describe a detour from tyrosine to coumarate (C00811). This last, suboptimal pathway cannot be found in the KEGG map for phenylpropanoid biosynthesis.

**Figure 4:**
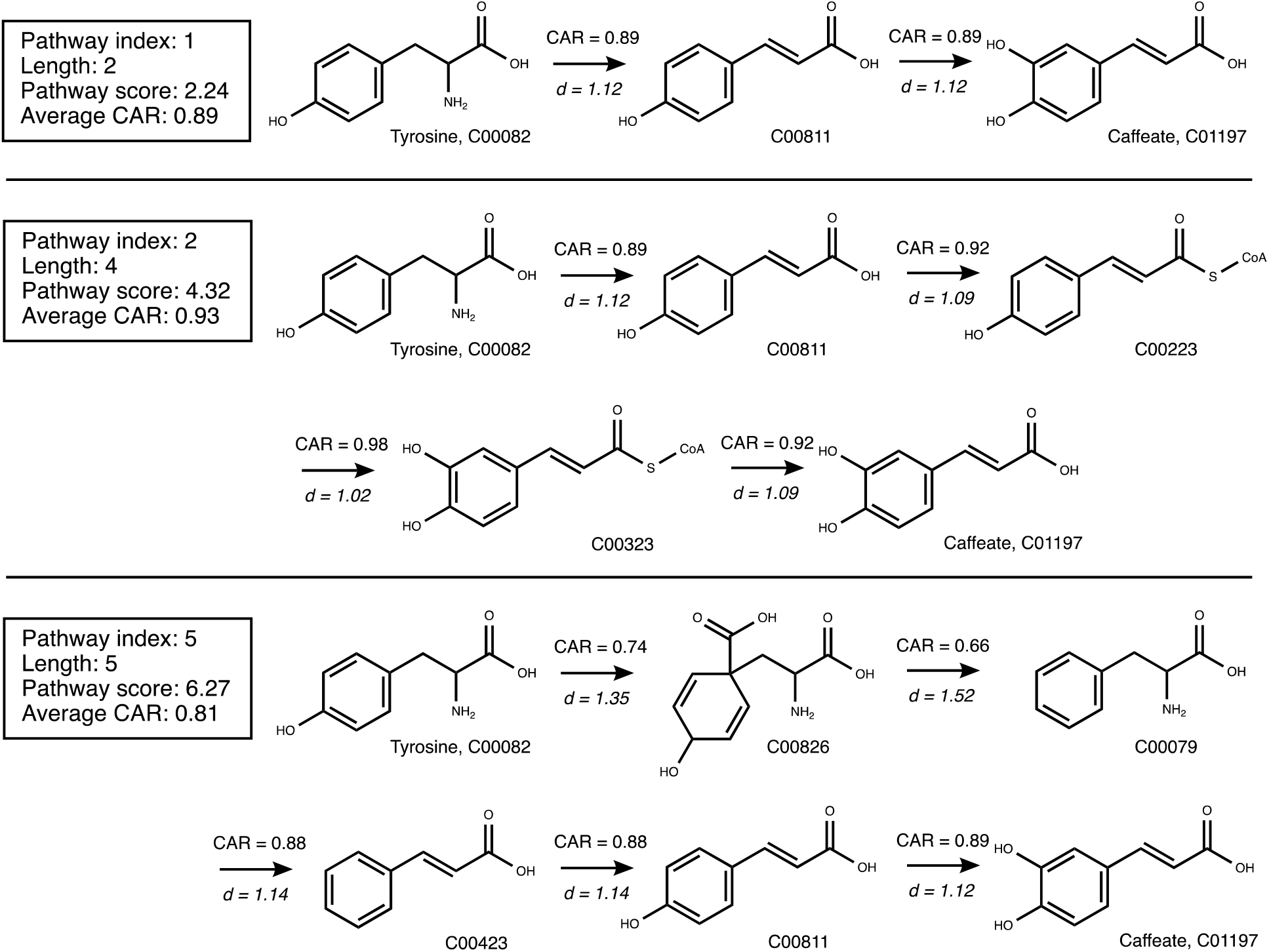
The pathways from Table 1 connecting tyrosine and caffeate with index numbers 1, 2, and 5 are visualized in detail for comparison. For each biotransformation, the CAR value as well as the default distance (d) are indicated.

In a second example, we searched for pathways connecting the compounds tyrosine and syringin. The number of pathways to be found was restricted to five, and we used three different transformations to calculate the distance between reactant-product pairs: The default transformation 1/CAR, the square root transformation, and the exponential transformation. Using the default option, NICEpath first listed three short pathways with a low average CAR (~0.5), followed by two longer pathways with high average CAR (~0.8) (Table 2). The square root option yielded only short pathways with a low average CAR, while the exponential option only resulted in longer pathways of high average CAR. Interestingly, all the long pathways with high CAR were identified as known metabolic pathways in KEGG, indicating that increasing the influence of the CAR on the distance by choosing an exponential transformation operator is helpful to reliably extract longer pathways.

**Table 2:**
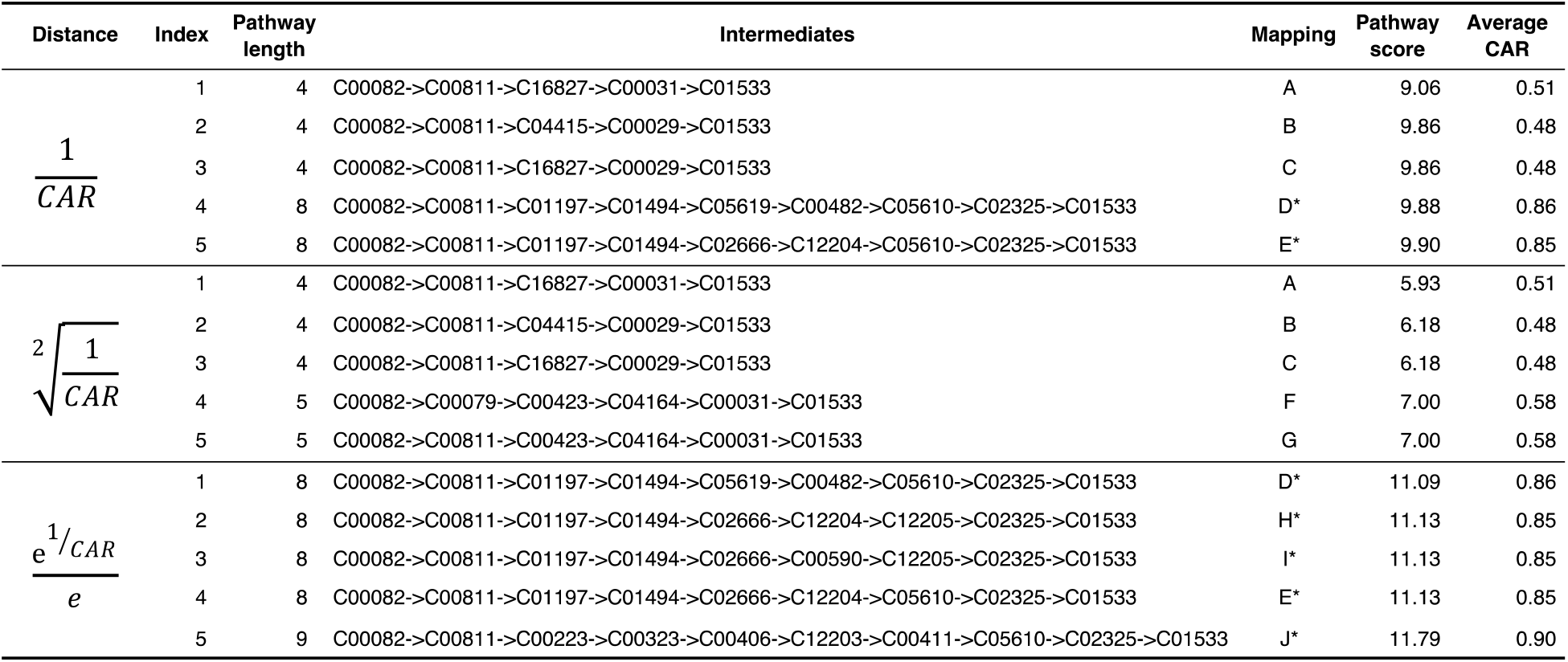
Output of pathway search from tyrosine (C00082) to syringin (C01197). Three different CAR transformations were used to calculate the distance between pairs of reactants and products: The first one (1/CAR) is the default option in NICEpath, the second is a square root transformation that decreases the effect of atom conservation, and the third one is an exponential transformation that increases the effect of conserved atoms in reactant-product pairs. Pathways are mapped across distance transformations with letters indicated in the column labeled “Mapping”. Pathways marked with an asterisk correspond to known metabolic pathways that fulfill the criteria for biologically relevant pathways.

Two pathways were chosen to understand in detail the influence of the type of transformation used for calculating the distances between reactants and products: one was short with a low CAR (A) and one was long with a high CAR (D*) (Figure 5). Pathway A connected tyrosine to syringin in four reaction steps, with a relatively low average CAR of 0.51. As already indicated by the low CAR, the pathway turned out to be a shortcut through glucose, with no atoms conserved between tyrosine and syringin. The pathway was ranked first in the default and the square root transformation types, but, interestingly, ranked 1114^th^ in the exponential case when re-running the search with an upper limit of 2000 pathways. The exponential transformation increases the penalty of atom loss in biotransformation, which leads to a higher pathway score assigned to the shortcut pathway. Pathway D* connected tyrosine to syringin in eight reaction steps, with a high average CAR of 0.86. It was ranked first using an exponential transformation, ranked fourth using the default distance calculation, and ranked 43^rd^ for the square root case. This second pathway kept the molecular core structure of tyrosine and modified it to produce syringin, conserving a maximum number of atoms. Remarkably, this pathway is part of the KEGG pathway map for phenylpropanoid biosynthesis, and it can therefore be called a confirmed, biologically meaningful pathway.

**Figure 5:**
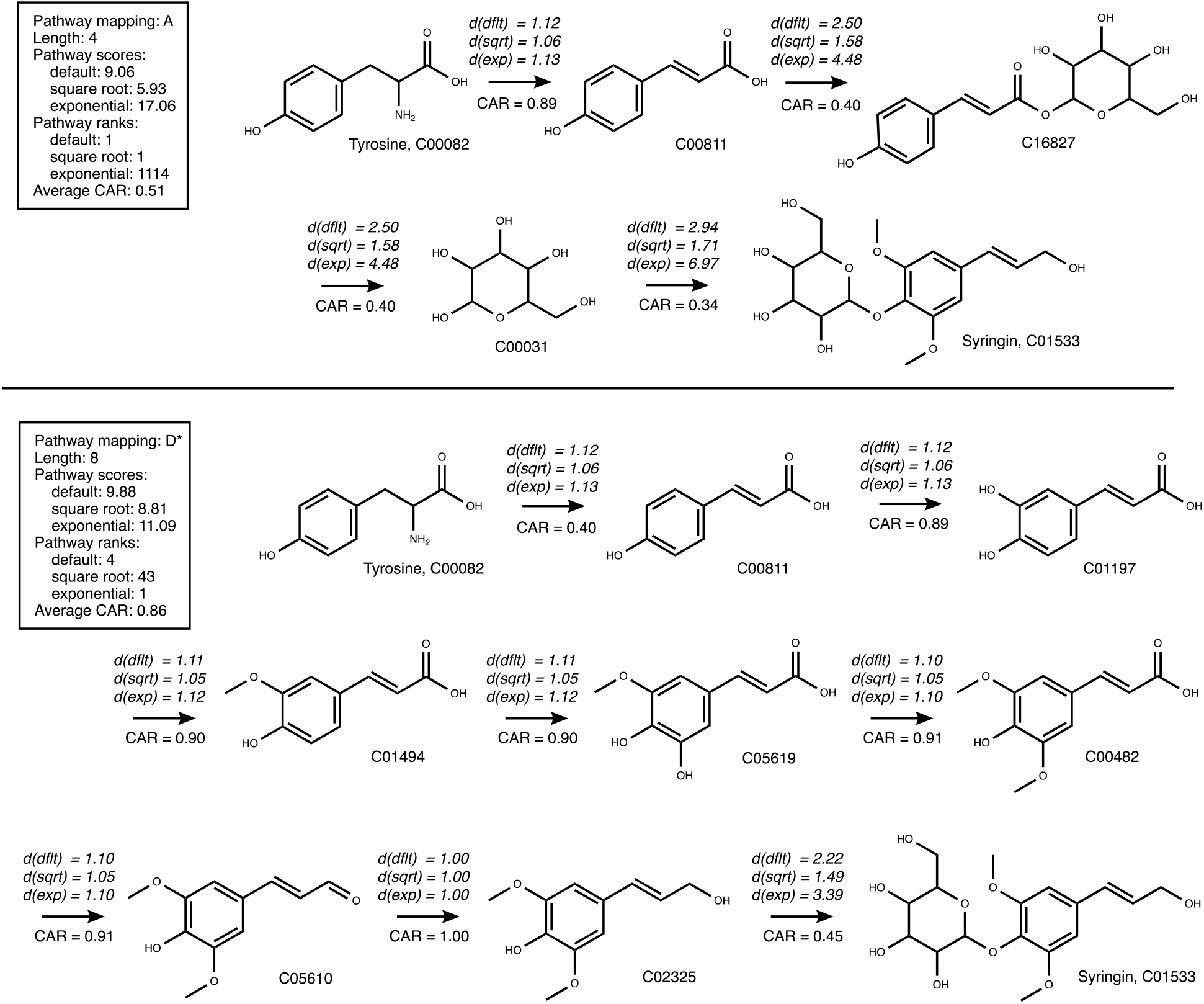
Comparison of two pathways (A and D*) from the pathway search connecting tyrosine to syringin. For each biotransformation, the CAR value along with the default distances for each transformation are indicated. (dflt): default distance, d(sqrt): square root transformation, d(exp): exponential transformation.

These two examples of pathway search problems illustrate the capacity of NICEpath to efficiently extract biologically relevant pathways from large biochemical networks. The algorithm robustly handled searches for long pathways of eight and more biotransformation steps, as they are usually present in secondary metabolism.

### Limitations and future challenges

There are cases in which NICEpath will not find satisfactory solutions. Possible reasons for suboptimal results are (i) the network does not contain the necessary reactions to connect the starting compound to the target compound, and (ii) the source and the target compound initially have only a few atoms in common. The first issue can be solved by adding the missing reactant-product pairs to the network. Missing steps can be hypothesized manually or predicted using reaction prediction tools such as BNICE.ch. The second issue is more complex, since it depends on the molecular structure of the source and target compound, as well as on the real number of biochemical transformations needed to transform one into the other. Possible solutions to improve the output include breaking down the search into several sub-searches by identifying intermediates and increasing the penalty on atom loss by using an exponential transformation of the CAR into the distance between reactants and products.

While our algorithm successfully circumvents the recurrent problem of shortcuts through small hub metabolites, it does not satisfactorily avoid shortcuts through big hub metabolites such as Coenzyme A (CoA). In fact, reactant pairs involving CoA structures on both sides have a lot of atoms in common, and therefore a high CAR value. For this reason, NICEpath excludes CoA by default from the reactant pair network.

## Conclusion

We introduce a new pathway search method based on weighted reactant-product pairs. To our best knowledge, this is the first to use automatically generated atom-weighted reactant-product pairs in combination with a k-shortest graph search approach. We benchmarked our method for reactant-pair weighting against the KEGG RPAIR database, and we discussed the advantages of NICEpath on practical examples. The strong point of NICEpath is that it is suitable for big biochemical networks, spanning more than hundreds of thousands biochemical reactions, such as hypothetical reaction networks generated by retrobiosynthesis tools and predictive biochemistry^29^. We estimate that the future development of reaction prediction tools, based on biochemical reaction rules or machine learning methods, will yield big hypothetical reaction networks that require optimized search tools to efficiently extract biochemical pathways. Furthermore, the presented method to translate metabolic networks into a graph structure can be used in the future to analyze the global characteristics of biochemical networks, such as the diameter of a network or its connectivity, and finally to detect and map knowledge gaps in metabolic databases.

Finally, the herein proposed framework will lay the foundation for further developments. Other types of weights, such as kinetic and thermodynamic considerations, can be integrated into the weighting of substrate-product pairs to steer the pathway search towards biochemically feasible pathways, and a set of user-defined parameters will make it easy to fine-tune the pathway search. The NICEpath code is available on GitHub (https://github.com/EPFL-LCSB/nicepath), and it comes with a collection of 5,434 known metabolic reactions with pre-calculated atom-weighted reactant pairs.

## Acknowledgements

We thank Kaycie Deyle for valuable feedback in the preparation of the manuscript. Funding for this work was provided by the Swiss National Science Foundation (SNSF) under Grant number 2013/158 (MicroScapeX) and the École Polytechnique Fédérale de Lausanne (ÉPFL).

## Author contributions

JH and VH designed the study. JH performed the implementation and the analyses, and wrote the manuscript.

